# Identifying the immunocompetence markers at weaning to predict long-term vaccine response and persistence in pigs

**DOI:** 10.64898/2026.10.13.758754

**Authors:** Abdullah Al Momen Sabuj, Marie Courbariaux, Catherine Denis, Gaëtan Lemonnier, Jordi Estellé, Marie-Hélène Pinard-van der Laan, Claire Rogel-Gaillard, Fany Blanc

## Abstract

Vaccination is a key strategy for controlling infectious disease in pigs, but the magnitude and persistence of vaccine-induced immunity vary among individuals. We investigated whether immune and physiological traits measured at weaning could predict long-term antibody (Ab) status and Ab persistence following vaccination against *Mycoplasma hyopneumoniae* (MH) and Influenza A virus (IAV), and whether predictive markers were shared between vaccines.

Two vaccination trials were conducted in a cohort of 381 Large White pigs (MH, n=189; IAV, n=192). Animals were vaccinated at weaning (28 days of age) and received a booster 21 days later. Long-term Ab status was defined at 118 days post-vaccination (dpv) using assay-specific thresholds, while Ab persistence was defined as the decline from the individual peak response at 28 or 35 dpv to 118 dpv. A broad panel of immune and physiological traits measured at weaning was evaluated as candidate predictors. Sparse Partial Least Squares Discriminant Analysis (sPLS-DA) was used for variable selection, followed by PLS-DA classification models.

Cross-validated accuracies ranged from 64.7% to 67.7%, indicating moderate predictive performance. Across vaccines, eight predictors were shared for long-term Ab status and twelve for Ab persistence. Hematocrit, IgG, platelet count, and plateletcrit (PCT) were associated with positive long-term Ab status for both vaccines, whereas mean corpuscular hemoglobin concentration (MCHC) and neutrophil percentage were associated with negative status. For persistence, higher vitamin D, α-1 globulin concentration, platelet count, and PCT were consistently associated with persistent responses across vaccines, while MCHC and spectral absorbance traits were associated with non-persistence.

These findings indicate that early-life immune and physiological traits measured at weaning are associated with immunocompetence traits that contribute to variation in subsequent long-term vaccine antibody outcomes and responsiveness. The identification of both shared and vaccine- or phenotype-specific predictors suggests that vaccine responsiveness reflects both general features of baseline immunocompetence and context-specific determinants.

## 1. Introduction

Respiratory diseases remain a persistent challenge in modern pig production, affecting animal health, welfare, and farm profitability (Saade et al., 2020; Boeters et al., 2023). Pathogens such as *Mycoplasma hyopneumoniae* (MH) and Influenza A virus (IAV) are particularly prevalent and are linked to reduced growth performance, increased mortality, and significant economic losses in commercial herds (Fano et al., 2007; Calderón Díaz et al., 2020). These pathogens play a major role in the porcine respiratory disease complex, a multifactorial condition that continues to challenge effective disease control strategies and highlights the need for sustained protective immunity throughout the production cycle (Fablet et al., 2012; Przyborowska et al., 2024).

Vaccination is widely used to mitigate the impact of respiratory infections in pigs and remains a central part of preventive health management (Rose and Andraud, 2017; Hoelzer et al., 2018). However, immune responses to vaccination vary markedly among individuals, even under standardized vaccination protocols (Adler et al., 2015; Zanella et al., 2015; Blanc et al., 2021, 2026). While some pigs develop strong immune responses that persist over time, others exhibit weak or short-lived responses, remain susceptible to infection, and may compromise overall herd immunity (Heininger et al., 2012; Blanc et al., 2021). Therefore, not only the magnitude but also the persistence of vaccine-induced immunity is critical for effective disease control (Chen et al., 2022). Animals that fail to mount or maintain protective responses may remain susceptible to infection and contribute to disease transmission at the herd level (Lipsit et al., 2022a). Although vaccine responses are often assessed shortly after vaccination, understanding the persistence of immunity over time is important because protection is expected to be maintained throughout the production cycle. In this context, the ability to mount and maintain vaccine-induced immunity reflects the broader biological concept of robustness, which describes an animal’s ability to maintain health and performance under challenging conditions (Shen-Orr and Furman, 2013; Friggens et al., 2017). Identifying animals at risk of poor or rapidly waning vaccine responses before vaccination could enable targeted management strategies, such as enhanced monitoring, revaccination, or isolation to protect herd health (te Beest et al., 2011; Knight-Jones et al., 2015; Lipsit et al., 2022a). Furthermore, understanding the biological determinants of vaccine response may improve vaccine design and application in heterogeneous populations (Pulendran et al., 2010; Kennedy et al., 2014).

Inter-individual variability in vaccine responsiveness is determined by host biology and is closely linked to immunocompetence, defined as the capacity to mount an appropriate immune response upon exposure to pathogens (Shen-Orr and Furman, 2013; Hine et al., 2014). Immunocompetence represents a multifaceted trait shaped by genetic background, physiological status, and environmental conditions, and can be assessed through measurable biomarkers (Mach et al., 2013; Llibre and Duffy, 2018; Jackson et al., 2020). In this context, blood serves as a valuable and accessible source of information, offering insights into both immune and metabolic status (Chaussabel, 2015). Biomarkers such as hematological parameters, immunoglobulin levels, plasma protein concentration, and natural antibodies measured in blood have been proposed as indicators of immunocompetence (Norris and Evans, 2000; Reid et al., 2021; Lipsit et al., 2022a). Variation in these biomarkers before vaccination is expected to contribute to differences in both the magnitude and persistence of vaccine-induced immune responses among individuals (Llibre and Duffy, 2018; Lipsit et al., 2020).

This early-life period surrounding weaning is particularly significant, as it represents a critical developmental stage marked by substantial physiological and environmental changes (Heo et al., 2013). Weaning involves separation from the sow, dietary shifts, social regrouping, and increased exposure to pathogens, all of which can affect immune development (Campbell et al., 2013; Tang et al., 2022). Concurrently, the decline in maternal antibodies increases reliance on the piglets’ own immune system protection (Bandrick et al., 2011).

Consequently, immune traits measured at weaning may reflect the immune system’s developmental status and provide valuable insights into subsequent vaccine-induced immune responses, as vaccination is often administered during this period.

Interest in vaccine-induced immunity is increasing. In pigs, host genetics, plasma cytokines, kinome profiles, and microbiota composition have all been associated with vaccine-induced antibody responses (Munyaka et al., 2019; Lipsit et al., 2020, 2022a). Similar associations have been observed in other species, reinforcing the value of early life biomarkers (Tsang et al., 2014; Berry et al., 2018; Jouneau et al., 2020; Gonçalves et al., 2021; Sevim Bayrak et al., 2024). However, these studies have generally focused on response magnitude measured shortly after vaccination, whereas relatively few have investigated predicting immune persistence over time or identifying predictive signatures shared across vaccines. In the same cohort of pigs vaccinated against MH and IAV, host genetics, blood transcriptome profiles, and gut microbiota composition have previously been associated with vaccine responsiveness (Munyaka et al., 2020; Blanc et al., 2021, 2026; Borey et al., 2021). However, researchers have not yet investigated easily measurable blood-based indicators of immunocompetence in this population. Because immune traits are highly interconnected, evaluating combinations of biomarkers may yield more robust predictors than considering individual traits in isolation.

We hypothesized that early-life immune traits measured at weaning are associated with vaccine-induced antibody responses, particularly their persistence, and that a subset of these biomarkers is shared across MH and IAV vaccine responses. Therefore, the study aimed to determine whether early-life immune traits can predict the persistence of vaccine-induced antibody responses in Large White pigs and to identify predictive signatures shared across MH and IAV vaccines.

## 2. Methods

### 2.1 Animal and experimental design

The animal experiment was conducted at the GenESI pig experimental facility of INRAE (Pig Innovative Breeding Experimental Facility; https://doi.org/10.15454/1.5572415481185847E12). The experimental population comprised Large White pigs from 48 families, produced in five successive batches. Sows (n = 47) were inseminated using semen from 48 boars selected to maximize genetic variability within the population. Within each litter, three to four piglets of each sex were selected based on body weight at 21 days of age to represent the average litter weight and avoid selection bias.

During the post-weaning phase (approximately 28 to 68 days of age), pigs were housed in pens of 20 to 30 animals and subsequently regrouped into pens of 10 to 12 animals during the finishing period (68 to 146 days of age). All animals received the same commercial diets throughout the experiment. Two vaccination trials were conducted within this population: one targeting *Mycoplasma hyopneumoniae* (MH, n=189) and the other targeting Influenza A virus (IAV, n=192). Detailed descriptions of the experimental populations and management conditions have been reported previously (Blanc et al., 2021, 2026). This study uses a subset of vaccinated animals from these cohorts to investigate predictors of vaccine-induced antibody persistence.

### 2.2 Vaccination and sampling schedule

Piglets were vaccinated at weaning (0 days post-vaccination, dpv), at an average age of 28 days (range: 24-31 days), followed by a booster vaccination at 21 dpv. In the MH trial, pigs received the commercial vaccine Stellamune (Pfizer), whereas animals in the IAV trial received the commercial trivalent vaccine Respiporc Flu3 (IDT Biologika). Blood samples were collected from the jugular vein at multiple time points to monitor the immune response. Serum samples were obtained at 0, 21, 28, 35, and 118 dpv to evaluate vaccine-induced antibody responses. Blood samples collected at weaning before vaccination were used to measure early-life immune parameters. Figure 1 presents the experimental design and the associated measures at each time point.

**Figure 1:**
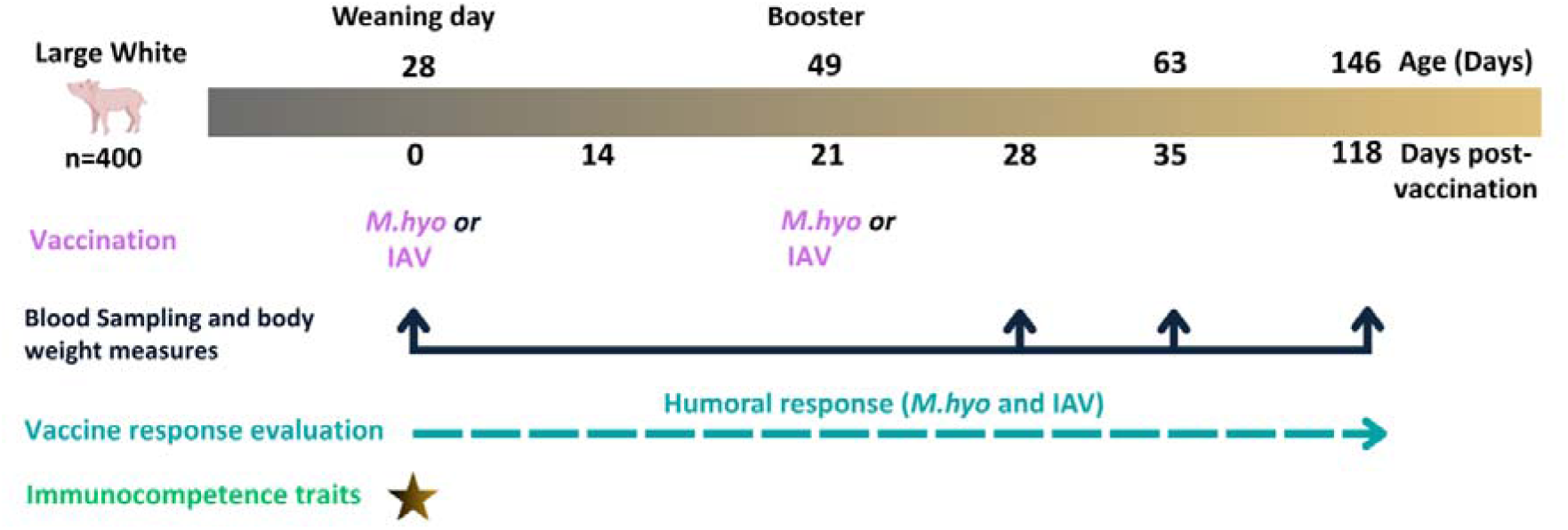
Experimental design of Large White pigs, showing vaccination, blood sampling time points and body-weight measurements, assessment of humoral responses to MH and IAV, and measurement of immunocompetence traits at weaning.

### 2.3 Measurement of vaccine-specific antibody responses

#### 2.3.1 MH antibody measurement

Serum antibodies specific to MH were quantified using a commercial enzyme-linked immunosorbent assay (ELISA) kit (IDEXX *M. hyopneumoniae* Ab test, IDEXX Europe B.V., The Netherlands) according to the manufacturer’s instructions, as previously described in Blanc et al. (2021). Serum samples collected at 0, 28, 35, and 118 dpv were analyzed in duplicate at a 1:40 dilution. Antibody levels were reported as sample-to-positive (S/P) ratios, calculated by dividing each sample’s corrected absorbance by that of the positive control, following the IDEXX procedure. According to the manufacturer’s thresholds, samples with an S/P ratio ≥0.4 were considered positive, values between 0.3 and 0.4 were classified as suspect, and an S/P <0.3 was considered negative. For subsequent analyses, samples classified as suspect (0.3–0.4) were grouped with negative samples and considered as negative.

#### 2.3.2 IAV antibody measurement

Antibody responses to IAV were measured using hemagglutination inhibition (HAI) assays, as described in Blanc et al. (2026). Functional antibody responses were evaluated using HAI assays performed according to standard procedures. Serum samples were treated with a receptor-destroying enzyme, heat-inactivated, and serially diluted before incubation with viral antigens and turkey red blood cells. HAI titres were defined as the reciprocal of the highest serum dilution that inhibited hemagglutination. Titers below the detection limit (< 5) were assigned a value of 2.5, and HAI titers≥ 40 were considered indicative of seroprotection.

### 2.4 Definition of phenotypes

#### 2.4.1 Long-term antibody status

For each vaccine, the long-term serological status was defined as the antibody titer measured at 118 dpv. Animals were classified into a binary category as seropositive or seronegative based on assay-specific threshold values. For MH, an S/P ratio ≥0.4 was considered positive, whereas values <0.4 were considered negative. For IAV, antibody titers≥40 were considered positive, while titers <40 were classified as negative.

To facilitate comparison of predictive models across vaccines and simplify result presentation, the two categories were referred to as positive (Pos) and negative (Neg) throughout the study.

#### 2.4.2 Antibody persistence

To construct binary persistence phenotypes for classification analyses, animals were categorized by the magnitude of antibody decline between peak response and 118 dpv. For each animal, the highest titer observed at either 28 or 35 dpv was identified as the peak response. The decline from this peak to 118 dpv was then calculated using either the percentage decrease (MH) or the absolute drop (IAV).

For MH, pigs showing a decrease of greater than 50% between the peak titer and 118 dpv were classified as non-persistent, whereas those with a decrease ≤50% were classified as persistent. For IAV, animals exhibiting a reduction of at least four titer dilutions (16-fold decrease in HAI titer) were classified as non-persistent. In contrast, animals with a smaller decline were considered persistent. Thresholds were selected to reflect biologically meaningful declines in antibody levels while maintaining sufficient group sizes for statistical analyses. This classification yielded a categorical phenotype that reflected the durability of the vaccine-induced antibody response.

### 2.5 Early life immune measurements

Hematological parameters were measured on EDTA whole blood using an Element HT5 veterinary hematology analyzer (Heska/Antech Diagnostics). Measurements included total and differential leukocyte counts, erythrocyte counts and associated red blood cell indices, and platelet counts and indices. The complete list of hematological traits, abbreviations, and units is provided in Table 1.

**Table 1:**
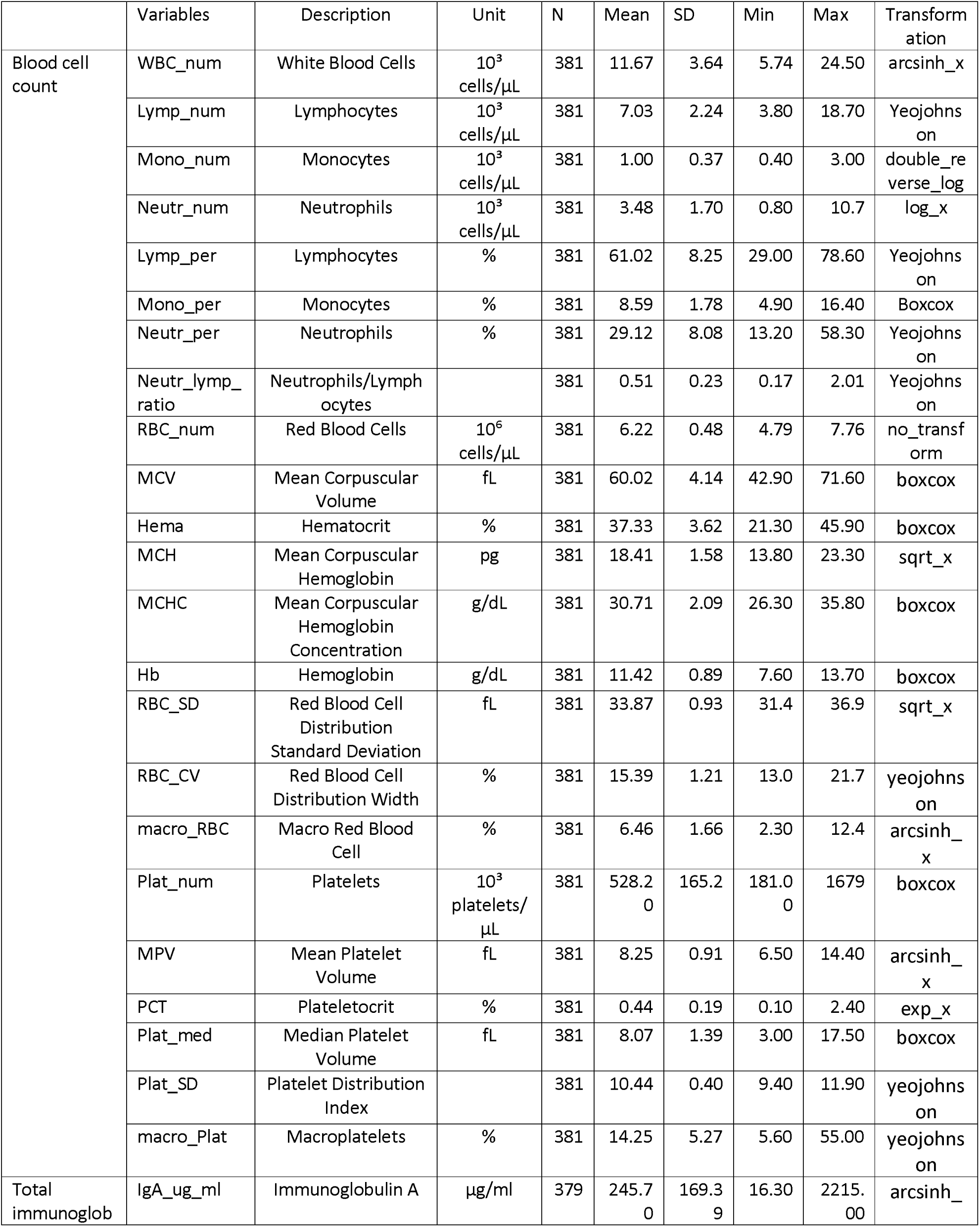
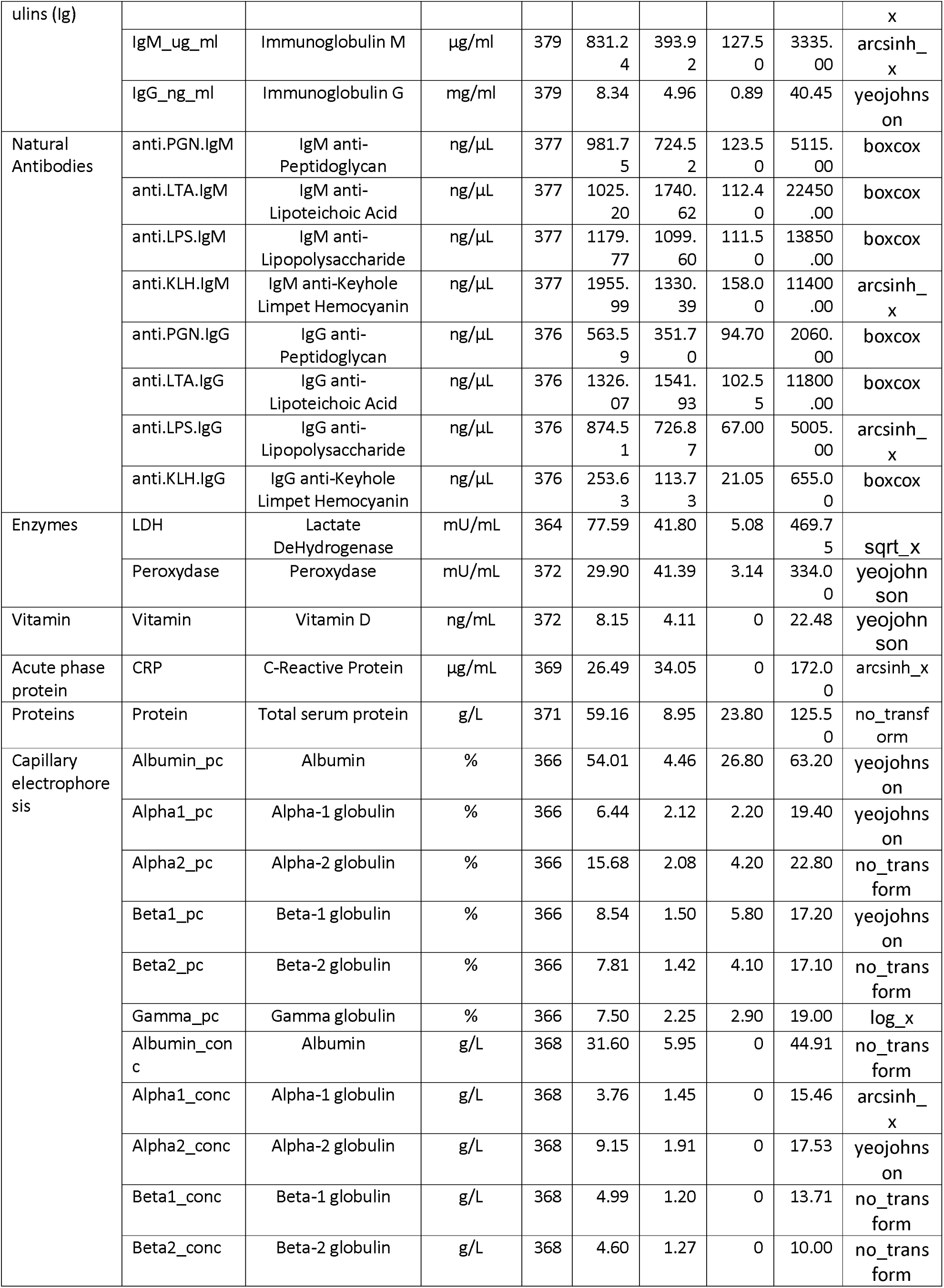
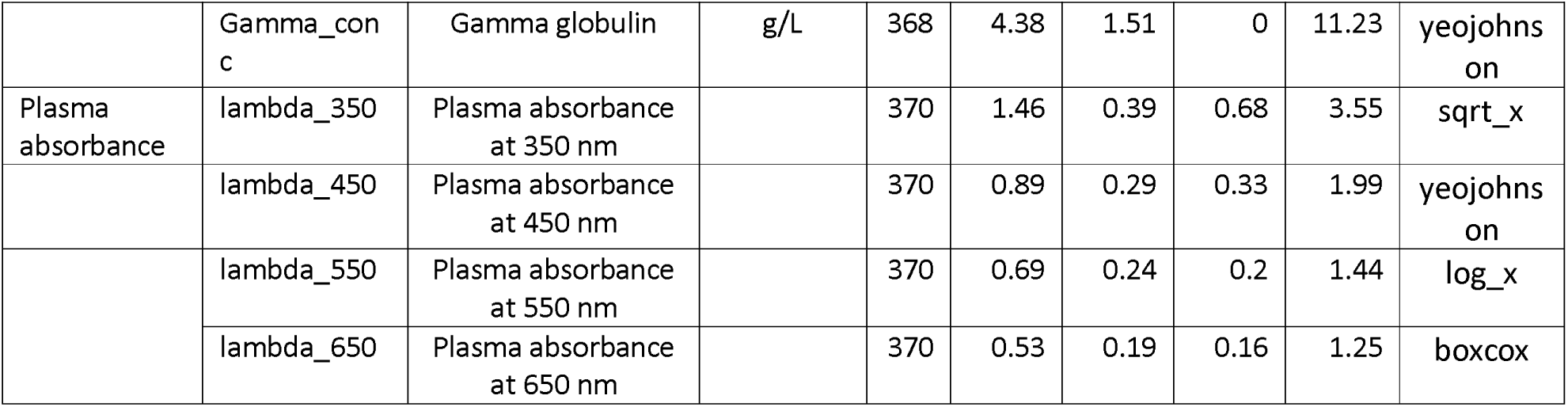
Descriptive statistics of immune parameters.

Plasma IgA, IgG, and IgM concentrations were measured by indirect ELISA using goat anti-pig IgA, IgG, or IgM capture antibodies and the corresponding HRP-conjugated detection antibodies (Bethyl Laboratories; IgA: BE-A100-102A/P, IgG: BE-A100-104A/P, IgM: BE-A100-117A/P). We determined concentrations from standard curves generated using Pig Reference Serum (Bethyl Laboratories). Absorbance was measured at 450 nm using a Multiskan FC microplate photometer (Thermo Fisher Scientific). All samples were analyzed in duplicate.

We measured natural antibodies specific to model antigens in plasma by indirect ELISA. Plates were coated with keyhole limpet hemocyanin (KLH; Megathura crenulata, Sigma H8283), lipopolysaccharide from *Escherichia coli* O55:B5 (LPS; Sigma L2880), lipoteichoic acid from *Bacillus subtilis* (LTA; Sigma L3265), or peptidoglycan from *Staphylococcus aureus* (PGN; Sigma 77140). Bound IgG and IgM natural antibodies were detected using the same HRP-conjugated goat anti-pig IgG and IgM antibodies as described above. A standard curve based on Pig Reference Serum was included on each plate, allowing semi-quantitative comparison of antigen-specific IgG and IgM levels across samples. All samples were analyzed in duplicate.

Total plasma protein concentration was measured using a bicinchoninic acid (BCA) colorimetric assay (Pierce™ BCA Protein Assay Kit, Thermo Fisher Scientific) according to the manufacturer’s instructions. Protein concentrations were calculated from a bovine serum albumin (BSA) standard curve. All samples were analyzed in duplicate.

Plasma protein fractions were determined by capillary electrophoresis using a MINICAP Protein 6 system (Sebia, France) according to the manufacturer’s instructions. Relative proportions (%) of albumin, α1-, α2-, β1-, β2-, and γ-globulin fractions were obtained from the electrophoretic profiles. Absolute concentrations (g/L) of each fraction were subsequently calculated using total plasma protein concentrations measured by the BCA assay.

Lactate dehydrogenase (LDH) activity was measured in plasma using a colorimetric assay kit (Lactate Dehydrogenase Activity Assay Kit, MAK066, Sigma-Aldrich) according to the manufacturer’s instructions. LDH activity was quantified from NADH production, and absorbance was measured at 450 nm. All samples were analyzed in duplicate.

Plasma peroxidase activity was measured in duplicate using a colorimetric assay with horseradish peroxidase (HRP) standards (Invitrogen) and TMB substrate (DY999, Bio-Techne). The reaction was stopped with 2 N sulfuric acid, and absorbance was measured at 450 nm. Activity was expressed in mU/mL.

Plasma C-reactive protein (CRP) concentrations were measured using a commercial porcine CRP ELISA kit (CRP-9, Life Diagnostics, USA) according to the manufacturer’s instructions.

Plasma 25-hydroxyvitamin D [25(OH)D] concentrations were measured in duplicate using the EUROIMMUN 25-OH Vitamin D ELISA kit (Ref. EQ 6411-9601, EUROIMMUN AG, Germany) according to the manufacturer’s instructions and expressed in ng/mL.

Plasma spectral absorbance traits were obtained by measuring the absorbance of 50 µL of undiluted plasma in half-area 96-well microplates (Greiner Bio-One). Absorbance was recorded at 350, 450, 550, and 650 nm using a Berthold Technologies microplate reader.

### 2.6 Statistical analysis

#### 2.6.1 Data Preprocessing

The initial datasets comprised 189 pigs vaccinated against MH and 192 pigs vaccinated against the IAV, each with 63 variables describing early-life immune traits and metadata. We combined the datasets to standardize preprocessing and data quality control. Samples with more than 20% missing values were excluded to minimize the potential bias from incomplete observations. We assessed variability in predictor variables to identify traits with near-zero variance and removed four variables. We imputed the remaining missing values using the missMDA R package, which performs multivariate imputation based on principal component analysis while preserving the correlation structure among variables. Multivariate diagnostic procedures identified potential outliers, leading to the removal of two samples. To improve comparability among predictors, we normalized all variables using the bestNormalize package, which selects the most appropriate normalizing transformation. The normality of transformed variables was evaluated using the Shapiro-Wilk test; detailed results are available in the supplementary material. Following preprocessing, the combined dataset was separated again by vaccine type for downstream analyses. After quality control, the baseline datasets comprised 185 MH-vaccinated and 184 IAV-vaccinated pigs and included 55 variables. For vaccine response phenotype, antibody measurements at 118dpv were available for 164 MH-vaccinated and 166 IAV-vaccinated pigs. These animals were subsequently classified based on long-term antibody status and persistence.

#### 2.6.2 Exploratory data analysis

Descriptive and exploratory analyses were performed to evaluate the distribution, structure, and interrelationships among early-life immune traits. Pairwise associations were quantified using Spearman correlation coefficients and visualized using the corplot function. Statistical significance was initially assessed at a nominal threshold of P < 0.05. To address multiple testing, p-values were adjusted using the False Discovery Rate (FDR) method via the p.adjust function in R. Only correlations that remained significant after FDR correction were considered robust.

#### 2.6.3 Sparse Partial Least Squares Discriminant Analysis

Sparse Partial Least Squares Discriminant Analysis (sPLS-DA) was conducted using the mixOmics R package (v6.10.8) to identify the most informative predictors of vaccine response. For each vaccine, we addressed two prediction tasks: classifying long-term antibody response (positive versus negative) and classifying antibody persistence (persistent versus non-persistent). Predictor variables included early-life immune markers measured at weaning as well as animal-related covariates, specifically sex, age at weaning, and weaning weight, which were incorporated as additional predictors in the models. Model tuning was performed using the tune.splsda function in mixOmics to determine the optimal number of components and variables retained per component. The tuning procedure was sequential, testing up to five components (ncomp = 5). For each component, the number of variables selected was evaluated within a candidate range of 5 to 58 predictors. Model performance was assessed using repeated M-fold cross-validation with five folds and 100 repetitions. The balanced error rate (BER) served as the primary performance metric to address potential class imbalance, and classification was based on the centroid distance. The number of predictors retained for each model was selected based on the BER profiles obtained during tuning, favoring parsimonious solutions at the point where further increases in the number of variables yielded only marginal improvements in classification performance. The tuning results indicated that a single component was sufficient to achieve the lowest classification error rates across the evaluated models. We then used predictor subsets selected from the BER profiles to construct the final classification models for both vaccine response phenotypes.

#### 2.6.4 Final PLS-DA models and model validation

We used the best predictors identified through the sPLS-DA tuning procedure to construct the final Partial Least Squares Discriminant Analysis (PLS-DA) classification models. We developed distinct models for each vaccine and phenotype. PLS-DA models were fitted using the plsda function, with the number of components determined by the tuning results. Predictor variables were standardized before model fitting. Model performance was evaluated using repeated five-fold cross-validation with 100 repetitions, implemented via the perf function. Classification accuracy was calculated from the balanced error rate using the following formula:

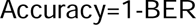

The discriminative performance of each model was further assessed by calculating the Area Under the Receiver Operating Characteristic Curve (AUC). We analyzed variable contributions to model components using loading plots to identify the predictors most strongly associated with classification outcomes.

#### 2.6.5 Identification of shared predictive markers

To determine whether similar predictors contributed to both response phenotypes, we compared variables selected in the long-term response and persistence models for each vaccine. We identified the intersection of selected predictors using set operations in R. We then compared predictors selected for MH and IAV to identify common markers across vaccines. Variables contributing to classification in both vaccine models were extracted by merging the corresponding loading tables with the dplyr package. We visualized overlaps among selected predictors using Venn diagrams with the ggVennDiagram package. We created separate diagrams for the long-term response and persistence models to illustrate shared and vaccine-specific predictive markers.

## 3. Results

### 3.1 Overview of immune parameters

We measured 55 immune and physiological traits at weaning. Table 1 provides detailed descriptive statistics and the transformations applied before modeling. Substantial inter-individual variation was observed across most traits. Hematological parameters showed broad distributions, particularly for leukocyte and platelet-related traits. Similarly, total immunoglobulins, natural antibodies, C-reactive protein (CRP), Lactate dehydrogenase (LDH) activity, and vitamin D concentrations exhibited marked variability among animals. Variability was also observed for electrophoretic protein fractions and plasma absorbance traits measured at different wavelengths.

### 3.2 Correlation structure of immune parameters

Spearman correlation analysis identified 628 significant pairwise correlations (*P* ≤ 0.05), with 498 correlations remaining significant following false discovery rate (FDR) correction. Hierarchical clustering identified several biologically meaningful trait groups (Figure 2).

**Figure 2:**
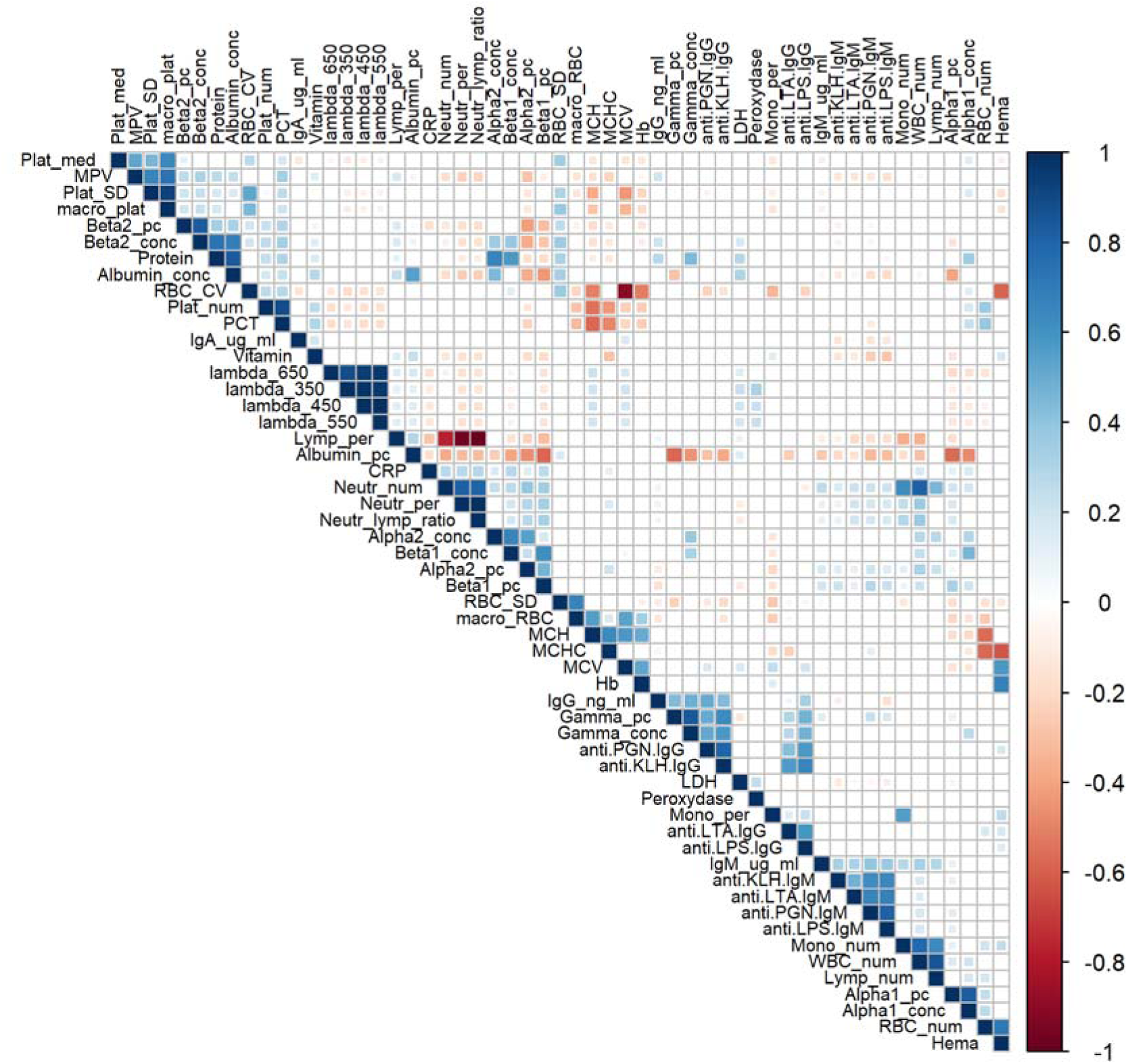
Spearman correlation matrix of baseline immunocompetence traits. Variables are ordered using complete-linkage hierarchical clustering. Square size and color intensity indicate correlation magnitude; blue denotes positive correlations, and red denotes negative correlations. Only statistically significant correlations (P ≤ 0.05) are displayed.

The first cluster comprised platelet-related traits (Platelet count, PCT, MPV, Platelet SD, macro platelets), which were positively correlated (r = 0.64-0.92). The second cluster included spectral absorbance measurements (Lambda 350-650 nm; r = 0.88-1.00). The third cluster grouped hematological traits, including RBC counts, Hematocrit, Hemoglobulin, MCV, MCH, and MCHC. Strong positive correlations were observed between Hematocrit and RBC counts (r = 0.72) and between Hemoglobin and Hematocrit (r = 0.67). In contrast, MCV was strongly negatively correlated with RBC CV (r = −0.90). Immune cell counts also formed a cluster, with WBC count strongly correlated with lymphocyte (r = 0.86), neutrophil (r = 0.82), and monocyte counts (r = 0.78). The strongest negative correlations were observed between % of Lymphocytes and % of Neutrophils (r = −0.96) and between % of Lymphocytes and Neutrophil lymphocytes ratio (r = −0.98). Another distinct cluster included natural antibodies and immunoglobulin-related traits, such as anti-PGN IgM and anti-LPS IgM (r = 0.82) and anti-KLH IgG and anti-PGN IgG (r = 0.81). Protein fraction traits also showed strong associations.

The observed correlation structure indicated redundancy within biological trait families and complementarity across immune compartments.

### 3.3 Antibody response profiles and classification

#### 3.3.1 Long-term antibody profiles and classification

Antibody responses against MH and IAV followed a similar temporal pattern, with titers increasing after the 21 dpv booster, peaking at either 28 or 35 dpv and declining by 118 dpv. The distribution of antibody responses at each sampling time, together with the seropositivity thresholds, is shown in Figure 3.

**Figure 3:**
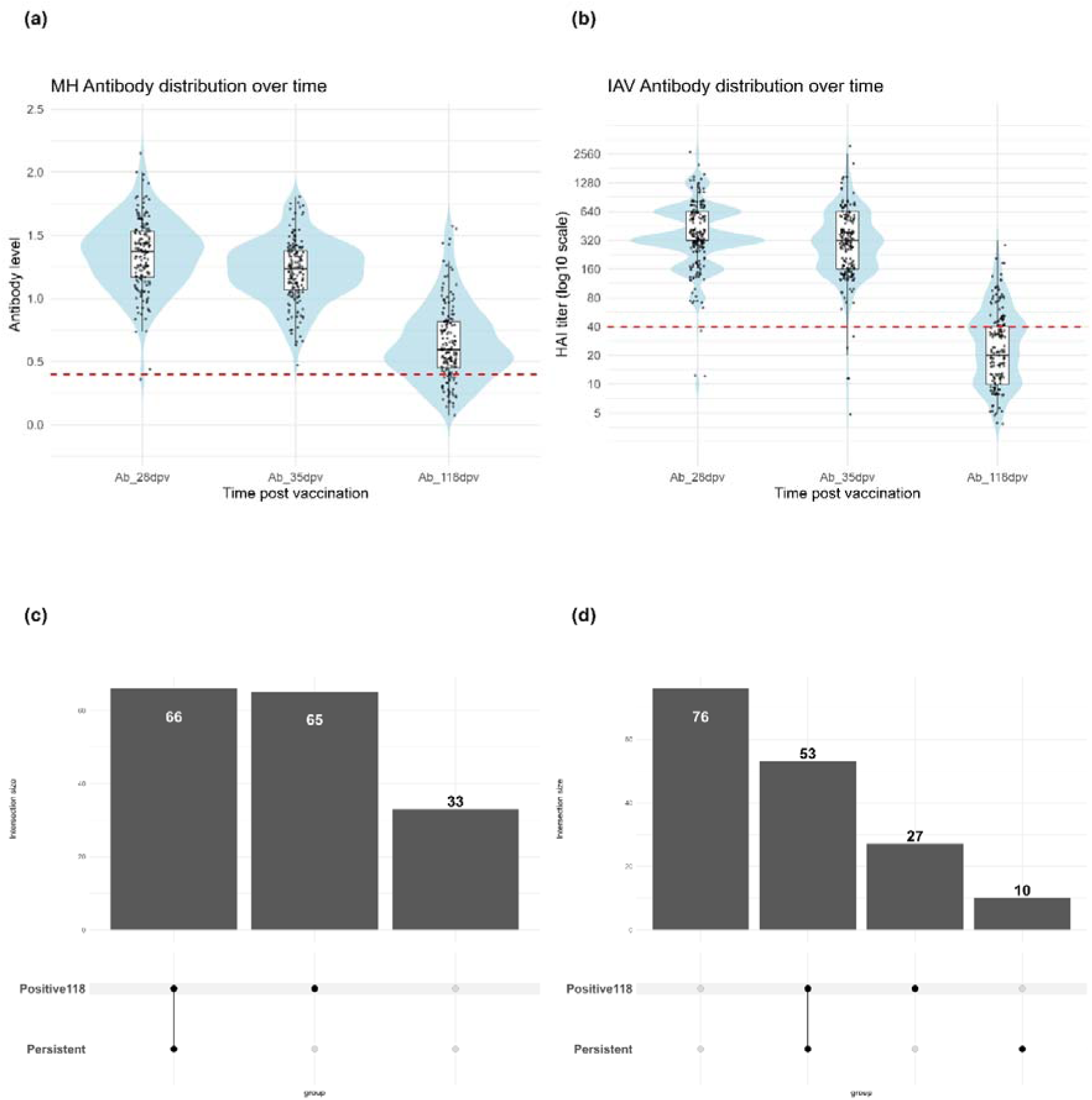
Antibody response profiles and classification of long-term Ab status and Ab persistence following vaccination. (a) Distribution of MH antibody levels at 28, 35, and 118 dpv; (b) Distribution of IAV HAI titers at the same time points, shown on a log10 scale. Red dashed lines indicate the predefined seropositivity thresholds for MH (S/P ratio ≥ 0.4) and IAV (HAI titers ≥ 40). (c) UpSet plot showing the overlap between long-term MH antibody status at 118 dpv and antibody persistence; (d) UpSet plot showing the corresponding overlap for IAV. The intersection sizes indicate the number of animals belonging to each combination of long-term Ab status and persistence.

At 118 dpv, animals were classified into two binary phenotypes according to vaccine-specific thresholds. For MH, 131 of 164 pigs (79.9%) were classified as positive and 33 (20.1%) as negative. For IAV, 80 of 166 pigs (48.2%) were classified as positive and 86 (51.8%) as negative at 118 dpv.

#### 3.3.2 Antibody persistence patterns

In MH, antibody titers declined at varying rates among animals. Using the predefined criterion of a greater than 50% decrease, 66 (40.2%) animals were classified as persistent and 98 (59.8%) as non-persistent.

For IAV, persistence classification relied on changes in HAI titres measured between the peak response and 118 days post-vaccination (dpv). Applying the threshold of a decrease of at least four titre dilutions, 63 (38.0%) animals were classified as persistent and 103 (62.0%) as non-persistent.

#### 3.3.3 Relationship between long-term antibody status and persistence

The UpSet plots illustrate the relationship between long-term antibody status and persistence (Figure 3c and 3d). For MH, all persistent animals were positive at 118 dpv, whereas the non-persistent group included both positive (n = 65) and negative (n = 33) animals.

For IAV, persistence was observed among positive animals (n = 53), while the positive group also included 27 non-persistent animals. In addition, 76 pigs were neither persistent nor positive, and 10 pigs were persistent despite not reaching the seroprotection threshold at 118 dpv.

### 3.4 Predictive performance of sPLS-DA and PLS-DA models

#### 3.4.1 MH vaccine

For the long-term Ab status, the sPLS-DA tuning procedure identified a single informative component and retained 20 predictors. The resulting PLS-DA model achieved a cross-validated accuracy of 67.7% and an AUC of 0.74.

Figure 4a shows the variables that most strongly discriminate between positive and negative Ab status at 118 dpv. Animals with a positive Ab status were primarily characterized by higher values of traits related to erythrocyte status (RBC count and hematocrit), humoral immunity (IgG), natural antibodies, and platelet-related traits (PCT and platelet count). In contrast, animals with a negative Ab status were characterized by higher values of MCHC, CRP, beta1 globulin fraction, neutrophil percentage, and weaning weight.

**Figure 4:**
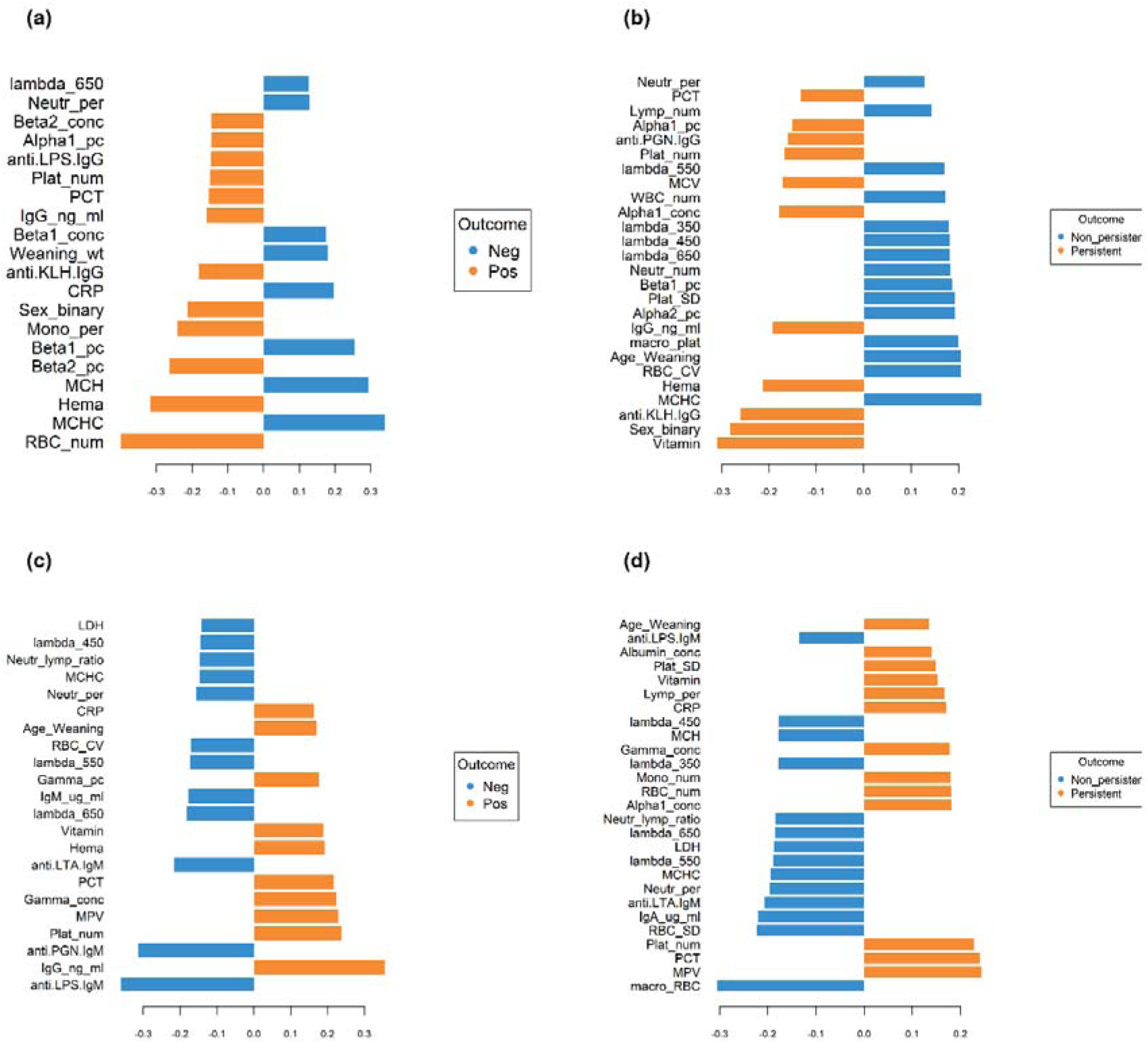
Immunocompetence traits loading on component 1 of the PLS-DA models for long-term Ab status and Ab persistence. Panels show MH long-term Ab status (a), MH antibody persistence (b), IAV long-term Ab status (c), and IAV antibody persistence (d). Positive and negative loadings indicate the direction of each variable’s contribution to the respective phenotype.

For antibody persistence classification, the tuning procedure retained 26 predictors and a single informative component. The corresponding PLS-DA model achieved a cross-validated accuracy of 66.6% and an AUC of 0.74.

Figure 4b shows the variables most strongly associated with persistence classification. Persistent animals were characterized by higher vitamin D concentrations, sex (higher proportion of persistent animals among females than males), hematocrit, anti-KLH IgG levels, humoral immunity (IgG), alpha 1 globulin concentration, platelet count, and PCT. Conversely, non-persistent animals were characterized by higher MCHC, beta1 globulin concentration, age at weaning, platelet distribution width, neutrophil count, and several spectral absorbance traits.

#### 3.4.2 IAV vaccine

For the long-term Ab status, the sPLS-DA tuning procedure identified a single informative component and retained 22 predictors. The resulting PLS-DA model achieved a cross-validated accuracy of 66.3% and an AUC of 0.71.

Figure 4c shows the variables most strongly associated with class discrimination. Animals with a positive Ab status at 118 dpv were primarily characterized by higher values of traits related to humoral immunity (IgG levels), platelet biology (platelet count, MPV, and PCT), erythrocyte status (hematocrit), CRP, and vitamin D concentration. In contrast, animals with a negative Ab status at 118 dpv were characterized by higher levels of total IgM, neutrophil percentage, neutrophil-lymphocyte ratio, natural antibodies against LPS, PGN, and LTA, as well as several plasma absorbance traits.

For antibody persistence, the tuning procedure retained a single informative component and 27 predictors. The resulting PLS-DA model achieved a cross-validated accuracy of 64.2% and an AUC of 0.67.

Figure 4d shows the variables that contribute most to persistence classification. Persistent animals were characterized by higher values of platelet-related traits (MPV, PCT, and platelet count), alpha-1 globulin concentration, CRP, and vitamin D levels. Conversely, non-persistent animals had higher proportions of macrocytic RBCs, elevated IgA and anti-LTA IgM levels, higher neutrophil percentages, and increased LDH activity.

### 3.5 Vaccine-specific predictors overlap between long-term Ab status and Ab persistence

#### 3.5.1 MH vaccine

For the MH vaccine, comparison of variables selected in the PLS-DA models for long-term Ab status and antibody persistence identified both shared and phenotype-specific predictors.

Eleven (31%) variables were common to both classification tasks, whereas 9 (26%) variables were unique to long-term Ab status and 15 (43%) were specific to persistence classification.

Figure 5a further details the shared variables and their directional contributions across both phenotypes. Several predictors related to humoral immunity (IgG and anti-KLH IgG), hematological status (hematocrit), platelet biology (platelet count and PCT), and alpha-1 globulins were consistently selected in both models.

**Figure 5:**
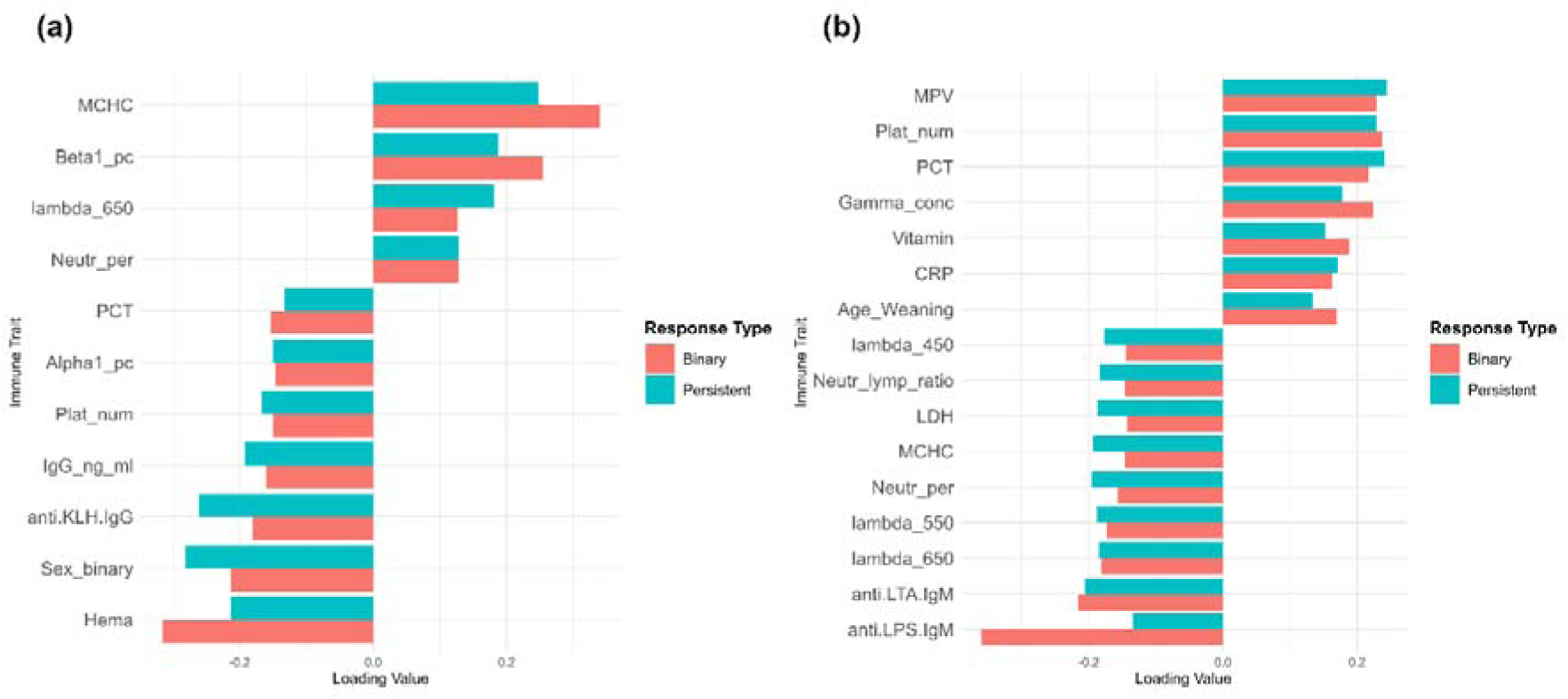
Shared predictors of long-term Ab status and Ab persistence. (a) PLS-DA loading values for variables shared between long-term Ab status and Ab persistence models for the MH vaccine; (b) Corresponding loading values for the IAV vaccine. Positive and negative loading values indicate the directions in which each variable contributes to the respective phenotype.

#### 3.5.2 IAV vaccine

For the IAV vaccine, a greater overlap between the two phenotypes. Of the selected variables, 16 (48%) variables were common to both the long-term Ab status and persistence models, whereas 6 (18%) were unique to the long-term response model and 11 (33%) to the persistence model.

Figure 5b summarizes the shared predictors and their contributions across both phenotypes. Platelet count, MPV, PCT, vitamin D levels, age at weaning, and CRP were consistently associated with positive long-term Ab status and persistent profiles. In contrast, anti-LPS IgM, anti-LTA IgM, neutrophil percentage, neutrophil-to-lymphocyte ratio, LDH, and spectral absorbance traits (Lambda_450, Lambda_550, and Lambda_650) were consistently associated with negative Ab status at 118 dpv and non-persistent profiles.

### 3.6 Shared and vaccine-specific predictors across vaccines

#### 3.6.1 Long-term Ab status

A comparison of predictors of long-term Ab status between the two vaccines revealed both shared and vaccine-specific variables. A total of 8 (24%) variables were common to both the MH and IAV models, while 12 (35%) were unique to MH and 14 (41%) were specific to IAV.

Figure 6a summarizes the shared predictors and highlights their directional contributions across both vaccines. Among these, hematocrit, IgG levels, PCT, and platelet count were consistently associated with a positive long-term antibody response in both the MH and IAV models. In contrast, MCHC and neutrophil percentage were consistently associated with a negative response in both vaccines. Notably, CRP showed opposite-directional associations between the MH and IAV models.

**Figure 6:**
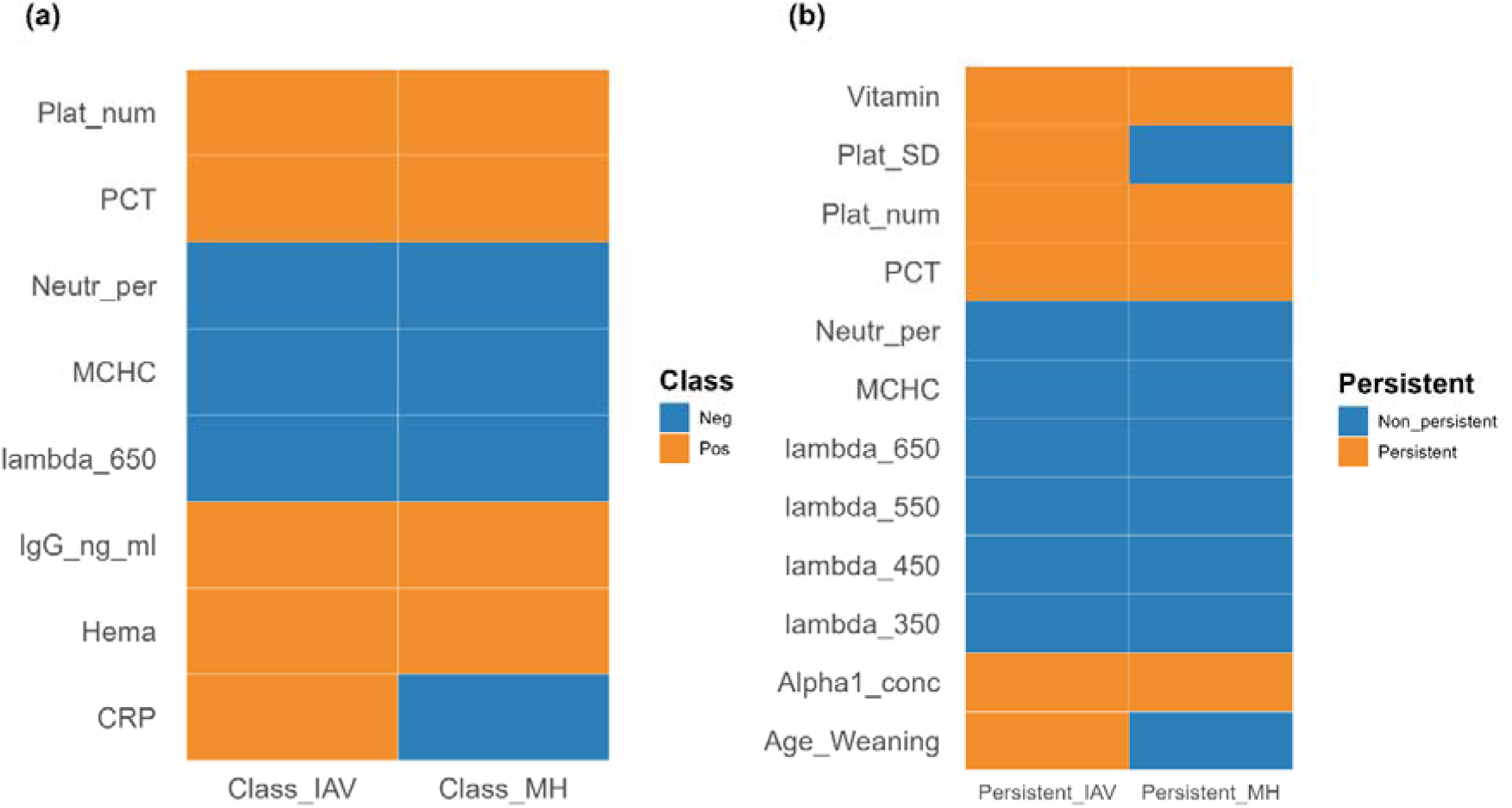
Shared predictors and their directional contributions across MH and IAV vaccine response models. (a) Shared predictors associated with positive or negative long-term Ab status at 118 dpv; (b) Shared predictors associated with persistent and non-persistent antibody responses.

#### 3.6.2 Antibody persistence

Regarding antibody persistence, there was a greater overlap of predictors between the vaccines. Of the selected variables, 12 (29%) were shared between the MH and IAV persistence models, while 14 (34%) were unique to MH and 15 (37%) were specific to IAV.

Figure 6b shows the shared predictors and their role in persistence classification for both vaccines. Variables such as vitamin D levels, alpha1_conc, platelet count, and PCT were consistently linked to persistent antibody responses in both MH and IAV. Conversely, MCHC, neutrophil percentage, and multiple spectral absorbance traits (lambda_350, lambda_450, lambda_550, lambda_650) were consistently connected with non-persistent responses.

Some predictors exhibited vaccine-specific patterns, including age at weaning and platelet distribution width (Platelet SD), which showed distinct associations with MH and IAV persistence models.

## 4. Discussion

### 4.1 Early-life immunocompetence biomarkers predict vaccine responses

This study demonstrates that early-life immune and physiological traits measured at weaning may predict antibody responses several months after vaccination. The predictive performance of the PLS-DA models was moderate, with accuracy ranging from 64% to 67% across both vaccines and phenotypes, indicating that early-life immune profiles explain part, but not all, of the inter-individual variation in vaccine responsiveness. This is consistent with the multifactorial nature of vaccine-induced immunity, which is influenced by baseline immunocompetence, genetic background, pathogen exposure history, nutritional status, stress, and environmental factors (Munyaka et al., 2019; Borey et al., 2021; Augustyniak and Pomorska-Mól, 2023). These results align with previous research in pigs showing that pre-vaccination host factors significantly influence vaccine outcomes. For example, Blanc et al. (2021) reported that pre-vaccination blood transcriptome profiles could distinguish high and low responders to MH vaccination, with functional enrichment in immune activation and cytokine signaling pathways differentiating the response groups. Similarly, Lipsit et al. (2020) identified kinome signatures in peripheral blood mononuclear cells prior to vaccination that predicted subsequent antibody responses, with pro-inflammatory cytokine signaling pathways dysregulated in low responders. Collectively, these studies and the present findings reinforce the concept that baseline immunological status before vaccination could be one of the determinants of vaccine antibody response.

### 4.2 Antibody persistence represents a distinct biological phenotype

Most of the studies of vaccine responsiveness evaluate immune responses at or near peak antibody levels, typically a few weeks after immunization (Lipsit et al., 2020, 2022b; Borey et al., 2021). However, the duration of the antibody response is also important, as animals are expected to maintain adequate immunity throughout the growing and finishing period.

Antibody persistence is increasingly recognized as a complementary aspect of vaccine performance, since peak antibody levels alone do not fully capture the duration of response (Chen et al., 2022; Palin et al., 2022). The substantial decline in antibody levels observed between the peak response at 28-35 dpv and 118 dpv for both MH and IAV indicates that a significant proportion of animals lose much of their initial response before reaching slaughter age. This finding demonstrates that antibody persistence represents a biologically and practically relevant phenotype, complementary to late antibody status. Nevertheless, persistence should be interpreted as a phenotype describing relative antibody decline, rather than as a direct measure of clinical protection, because antibody thresholds and the relationship between antibody titres and protection may vary between vaccines.

### 4.3 Common and specific predictors of vaccine responsiveness across vaccines

An objective of this study was to determine whether predictors of vaccine responsiveness are shared between two different vaccines and to evaluate whether these commonalities differed between long-term Ab status and Ab persistence. Eight predictors were identified as shared between vaccines for long-term Ab status, while 12 were shared for Ab persistence. These shared predictors suggest that certain relatively accessible blood-based traits may reflect underlying immune and physiological states that contribute to vaccine responsiveness across different antigenic challenges (Flori et al., 2011; Lipsit et al., 2022b). In addition, vaccine-specific predictors were also identified, indicating that some determinants of vaccine response are dependent on the pathogen and the vaccine nature (Van Tilbeurgh et al., 2021; Hagan et al., 2022). Consequently, the shared predictors should not be regarded as universal markers of vaccine responsiveness. Rather, they may represent components of broader baseline immunocompetence relevant across different immune challenges. This interpretation is particularly relevant given that the vaccine used in this study targeted biologically distinct pathogens and was evaluated using different antibody assays.

The patterns across four vaccine models indicate that the biological interpretation of the selected predictors depends on both the specific vaccine and the response phenotype. Platelet count and PCT were shared across vaccines for both long-term Ab status and Ab persistence, while MCHC and neutrophil percentage were associated with less favorable outcomes. In contrast, hematocrit and IgG were shared across vaccines, specifically for long-term Ab status, whereas vitamin D and α1-globulin concentration were shared specifically for persistence. Other predictors showed more vaccine-specific patterns.

#### 4.3.1 Platelet-related traits

Platelet count and PCT showed the most consistent associations across models, being associated with positive long-term Ab status and persistence for both vaccines. In contrast, MPV was selected only in the IAV models, indicating that this consistency did not extend to all platelet-related traits.

The repeated selection of platelet count and PCT across both vaccines and response phenotypes suggests that these traits may capture aspects of the host physiological or immune state at weaning that are relevant to both long-term Ab status, the magnitude, and antibody persistence/maintenance. Beyond their established roles in hemostasis, platelets interact with leukocytes and other immune cells and participate in inflammatory and immune regulatory processes through receptor-mediated signaling and release of soluble mediators (Morrell et al., 2014; Koupenova et al., 2018). Interestingly, previous transcriptomic analyses conducted in the same experimental population identified pre-vaccination transcriptional signatures involving platelet activation that were associated with later antibody response to *M. hyopneumoniae* vaccination (Blanc et al., 2021). Human vaccine studies have also linked platelet-related signatures to antibody durability and post-vaccination antibody kinetics (Flego et al., 2022; Cortese et al., 2025). However, these observations do not establish a direct functional relationship between baseline platelet traits and subsequent vaccine responses.

#### 4.3.2 Markers of humoral immune status

Humoral markers exhibited more phenotype-and vaccine-specific patterns than platelet-related traits. Total IgG was a shared predictor of positive long-term Ab status for both MH and IAV, but not of Ab persistence across vaccines, suggesting that the humoral environment before vaccination may be more closely associated with the later Ab level than with its maintenance after the peak. In contrast, gamma globulin and several natural antibody traits were selected only in specific vaccine or phenotype models.

At weaning, pigs undergo major significant changes in humoral immunity, antibody production, and immune development, with declining maternally derived antibodies and increasing endogenous antibody production, endogenous antibody production (Sinkora et al., 2002; Rooke et al., 2003). Baseline IgG at this age may therefore reflect the ongoing maturation and overall status of the humoral immune system rather than a specific determinant of subsequent vaccine responsiveness.

#### 4.3.3 Hematological traits

Hematological traits also showed consistent associations with vaccine outcomes. Hematocrit was positively associated with long-term Ab status for both vaccines and with persistence for MH, whereas MCHC was consistently associated with negative long-term Ab status and non-persistence across both vaccines. As these traits primarily reflect erythrocyte physiology, their associations may capture broader physiological differences among piglets at weaning rather than direct effects on vaccine responsiveness. The mechanisms underlying these associations remain unclear and warrant further investigation.

Neutrophil-related traits were also associated with less favorable vaccine outcomes. Higher neutrophil percentage was associated with negative long-term Ab status and non-persistence for both vaccines, and an elevated neutrophil-to-lymphocyte ratio was also selected in the IAV models. These traits can reflect inflammatory or physiological stress status, although neither serves as a specific marker of inflammation (Zahorec, 2021).

Because weaning itself includes substantial changes in circulating immune cell populations (Kick et al., 2011), these associations may partly reflect the physiological and immune state of the animals during this transition rather than specific determinants of vaccine responsiveness.

#### 4.3.4 Vitamin D and inflammatory markers

Vitamin D was a common predictor of antibody persistence for both vaccines, with higher concentration at weaning associated with persistent responses. It was also selected in the long-term Ab status in the IAV model, but not in the MH model, suggesting a particularly consistent association with the persistence phenotype. Vitamin D is involved in several aspects of immune regulation, including innate immunity, antigen presentation, and T- and B-cell differentiation (Sadarangani et al., 2015). However, associations between vitamin D status and vaccine-induced immunity have been inconsistent across vaccines and populations (Sadarangani et al., 2015; Lee et al., 2018). Thus, vitamin D status at weaning may represent a marker of the physiological and immunological state associated with subsequent Ab persistence, although the present study cannot establish a causal relationship.

Inflammatory markers showed less consistent patterns across vaccines and phenotypes. Higher α1-globulin concentration was associated with antibody persistence for both vaccines, whereas CRP was selected only for long-term Ab status and showed opposite directions in the MH and IAV models. CRP is a major acute-phase protein in pigs and is strongly influenced by inflammatory conditions (Petersen et al., 2004). Its opposite associations with MH and IAV vaccine outcomes suggest that the relationship between baseline inflammatory status and subsequent vaccine response may be context-dependent and may vary with the pathogen, vaccine formulations, baseline health state, or interaction with other predictors. The α1-globulin fraction includes proteins involved in the acute-phase response, including α1-acid glycoprotein, which has been shown to behave as a negative acute-phase protein in pigs (Heegaard et al., 2013). However, because the α1-globulin fraction was measured as a whole, its association with antibody persistence cannot be attributed to a specific protein or inflammatory mechanism.

#### 4.3.5 Natural antibodies and spectral absorbance traits

Natural antibody traits showed strongly vaccine-specific patterns. For IAV, higher IgM natural antibodies against LPS, LTA and PGN were associated with negative long-term Ab status, while anti-LPS and anti-LTA IgM were also associated with non-persistence. In contrast, higher anti-KLH IgG was associated with positive long-term Ab status and persistence for MH. Natural antibodies are part of the pre-existing humoral repertoire and may therefore provide information on the baseline immune state (Maddur et al., 2020). In pigs, natural antibodies against the same four antigens were heritable and showed genetic associations with resilience to a later disease challenge, although phenotypic correlations were weak (Chen et al., 2020), while associations with health in dairy cattle varied according to isotype and sample type (Thompson-Crispi et al., 2013). Thus, the contrasting associations observed here are more consistent with natural antibodies reflecting context-dependent differences in baseline immune status than with a uniformly favorable or unfavorable immune phenotype.

Several spectral absorbance traits, particularly wavelengths around 450-650 nm, were repeatedly associated with non-persistence and, for IAV, with negative long-term Ab status. These traits were highly correlated (r = 0.88-1.00), indicating that they likely capture a common feature of plasma composition rather than independent biological signals. In poultry, plasma absorbance around 450–490 nm overlaps with the absorption spectrum of circulating carotenoids and is sensitive to physiological changes induced by coccidial infection (Hamzic et al., 2015). Carotenoids have antioxidant and immunomodulatory properties and have been associated with both cellular and humoral immune responses (Chew and Park, 2004). Thus, variation in carotenoid status could contribute to the association observed at 450 nm. However, whether plasma absorbance provides a similar proxy for carotenoids in pigs remains to be established, particularly at the longer wavelengths measured here. Targeted biochemical analyses would be required to identify the plasma components underlying this spectral signature.

#### 4.3.6 Animal-related covariates

Animal-related covariates were selected in some models, but their associations varied according to vaccine and response phenotypes. Sex was selected in both MH models, with females more frequently associated with positive long-term Ab status and persistence, consistent with previous reports of sex-related differences in immune responses in pigs (Fardisi et al., 2023). Weaning age was selected as a predictor of persistence for both vaccines, but in opposite directions, with older age associated with persistence for IAV and non-persistence for MH. Additionally, it was also associated with positive long-term Ab status for IAV. Given the narrow weaning age (24-31days), and the contrasting associations between vaccines, these effects should be interpreted cautiously and may reflect other early-life physiological differences rather than direct effects of age on vaccine responsiveness.

### 4.4 Implication, limitation and future perspectives

The primary practical interest of these findings is that relatively simple blood-based traits measured at weaning provide information about antibody outcomes several months later. In comparison to transcriptomic, kinome, or microbiota signatures previously associated with vaccine responses in pigs, immunological and physiological phenotypes may offer greater accessibility for future immunocompetence phenotyping (Munyaka et al., 2019; Lipsit et al., 2020; Blanc et al., 2021; Borey et al., 2021). However, the current models demonstrate only moderate predictive performance and lack external validation. Consequently, these traits are not yet suitable for individual-level management or breeding decisions.

This study has several limitations. First, it was conducted in a single Large White experimental population under controlled conditions, which limits the generalizability of the findings to other breeds, farms, vaccination protocols, and production environments.

Second, the classification of persistence was based on vaccine-specific thresholds for antibody decline. Although these thresholds were chosen to represent substantial declines while maintaining adequate group sizes, alternative thresholds or continuous measures of antibody decay could yield different results. Third, the outcomes assessed were antibody-based phenotypes rather than direct measures of clinical protection, disease severity, or transmission, and did not capture cellular or mucosal immune responses. Fourth, correlations among several predictors indicate that selected variables should not necessarily be interpreted as independent biological signals. Finally, the use of internal repeated cross-validation provides an estimate of model performance within this study but does not substitute for external validation in independent populations.

Future research should therefore prioritize validation of these multivariate signatures in independent pig populations and, where possible, across different production environments and vaccines. Modeling antibody levels and decline trajectories as continuous phenotypes would help assess the robustness of the associations identified using binary persistence classes. Importantly, these candidate immunocompetence traits should be evaluated against relevant health and protection outcomes to determine the biological and practical significance. Integration with molecular and genetic data could subsequently help clarify the mechanism and genetic basis underlying these associations.

## Conclusion

In conclusion, this study demonstrates that early-life immune and physiological traits measured at weaning contain information on both long-term Ab status and Ab persistence following vaccination against MH and IAV in pigs. While some predictors, particularly platelet-related traits, markers of humoral immunity, and hematological traits, showed consistent associations across vaccines and response phenotypes, others were more specific to the vaccine or antibody phenotype considered. These findings support the potential of early-life immune profiling to characterize inter-individual variation in vaccine responsiveness and identify candidate indicators of immunocompetence. Validation in independent populations and against relevant health and protection outcomes will be required to determine their broader biological and practical relevance.

